# Sequence length controls coil-to-globule transition in elastin-like polypeptides

**DOI:** 10.1101/2023.12.04.569864

**Authors:** Tatiana I. Morozova, Nicolás A. García, Jean-Louis Barrat

## Abstract

Phase separation of disordered proteins resulting in the formation of biocondensates has received significant attention due to its fundamental role in cellular organization and functioning and is sought after in many applications. For instance, the liquid-liquid phase separation of tropoelastin initiates the hierarchical assembly process of elastic fibers, which are key components of the extracellular matrix providing resilience and elasticity to biological tissues. Inspired by the hydrophobic domains of tropoelastin, elastin-like polypeptides (ELPs) were derived which exhibit a similar phase behavior. Even though, it appeared almost certain that elastin condensates retain liquid-like properties, a recent experimental study questioned this viewpoint by demonstrating that the aggregate state of elastin-derived materials can depend on the length of hydrophobic domains. Here, we employ state-of-the-art atomistic modeling to resolve the conformational ensembles of a single ELP as a function of its sequence length in the temperature range relevant to possible applications. For the first time, we report the free energy profiles of ELPs in the vicinity of conformational transitions which show more compact polypeptide conformations at higher temperatures in accord with their thermoresponsive nature. We access the conformations visited by ELPs through descriptors from polymer physics. We find that short ELPs always remain in coil-like conformations, while the longer ones prefer globule states. The former engages in intrapeptide hydrogen bonds temporarily retaining their liquid-like properties while the latter forms long-lived (hundreds of nanoseconds) intra-peptide hydrogen bonds attributed to ordered secondary structure motifs such as *β*-bridges and turns. Our work demonstrates the importance of the sequence length as a modulator of conformational properties at a single chain and possibly explains the change in aggregate state in elastin condensates.

---

Phase separation of biological macromolecules resulting in their compartmentalization in distinct phases has been extensively realized both in nature and industrial applications^1,2^. In particular, intrinsically disordered proteins or regions (IDPs/IDRs) that possess high conformational heterogeneity^3^ are often involved in the formation of biomolecular condensates in cells^4^. One example is stress granules, which assemble as a response to cellular stress from RNA and RNA-binding proteins in the cytoplasm and are vital for the translational and signaling functions of cells^5^. Similarly, in the extracellular matrix, the coacervation of tropoelastin sets the stage for the multi-step assembly process of elastic fibres^6^, which provide resilience and elasticity to biological tissues such as lungs, blood vessels, skin^7^. These properties are encoded in the sequence of tropoelastin, which comprises alternating cross-linking and hydrophobic domains. The former is responsible for the durability of elastic fibers, while the latter drives self-assembling pathways^8^. Inspired by elastin, synthetic macromolecules such as elastin-like polypeptides (ELPs) resembling only hydrophobic repeats were derived^9^. ELPs mimic the ability of tropoelastin to undergo liquid-liquid phase separation characterised by a lower critical solution temperature (LCST) phase behavior in aqueous environments^9^. Typically, the critical temperature occurs near room temperature which gives rise to a number of possible applications including protein purification^10^, drug delivery^11^, and tissue engineering^12^.

While the formation of biomolecular condensates certainly plays a crucial role in organism functioning, deregulation in self-assembly processes and/or change in condensate properties are linked to neurodegenerative diseases, and cancer^4,13^. Thus, numerous works were dedicated to connecting sequence-conformational ensemble-function relations for IDPs and IDRs. In this context, predominately IDRs and IDPs that contain charged residues were investigated extensively as they make up a large fraction of the disordered proteome^14,15^. Order parameters based on the distribution of electric charge were developed that can predict sequence preference to sample extended or compact states^16–19^. By drawing the analogy between intra- and inter-protein interactions, sequences that visit compact conformations are more prone to phase separation, connecting individual and collective properties^20–22^. Naturally, these approaches do not apply to hydrophobic sequences, such as ELPs. Moreover, other system variations than sequence composition, also known to influence conformational properties of macromolecules, such as the chain length, have not been considered.

Dense, phase-separated state of elastin remains controversial. Earlier theoretical works suggested that elastin adopts compact conformations rich in *β*-turns formed by its hydrophobic side chain groups^23^. In contrast, others proposed models with high conformational entropy^24^. More recently, numerical works on the self-assembly of short ELPs (35 residues in length) showed the formation of well-hydrated aggregates in which individual chains adopt coil conformations with sparse and transient *β*-turns highlighting the dynamical, liquid-like nature of ELPs^25^. Similar results were later obtained for aggregates made of longer ELPs^26^. These numerical studies are in accord with the recent experimental work, which reports the formation of transient *β*-turns in highly dynamic hydrophobic domains resolved by NMR spectroscopy^27^. However, Ceballos and coworkers demonstrated experimentally that the aggregate state of elastin condensates can depend on the length of its hydrophobic domains, where the transition from liquid-like droplets to more solid-like occurs with increasing chain length of hydrophobic domains^28^. This work highlighted the importance of the chain length as a modulator of the material state of elastin condensates. However, the microscopic understanding of the observed experimental difference was difficult to deduce.

Here, we study the conformational and dynamical properties of a single ELP of the sequence GVG(VPGVG)_*n*_ as a function of the number of pentamers *n* in the temperature range that includes the critical temperature and relevant for possible ELPs applications. We employ state-of-the-art atomistic modeling in conjunction with advanced sampling techniques resulting in the total simulation time exceeding 40 microseconds. First, we reconstruct the free energy profiles of ELPs in the vicinity of the conformational transition, finding that more compact states are populated at higher temperatures at all chain lengths considered, consistent with a LCST behavior. We evaluate conformational ensembles visited by ELPs using descriptors adopted from polymer physics. Surprisingly, we find that short sequences with less than five pentamers always remain in coil-like states, in contrast to longer sequences which populate globule-like conformations. We rationalize our findings by computing the internal hydrogen bond formed between amino acids, their distribution, and timescales. We observe that long sequences (>60 residues) are capable of forming a few long-lasting hydrogen bonds (hundreds of nanoseconds) while the short ones engage in short lived interactions that last just over tens of nanoseconds demonstrating their dynamical nature. Even though the types of the secondary structure formed by all ELPs considered remain the same, their population differs significantly as a function of the chain length. From mostly coil elements (≈70 % for short ELPs) toward more structured ones with an increased number of hydrogen-bonded turns, bends, and *β*- bridges formed. The latter can exhibit remarkable longevity with hydrogen bonds lasting in total over 900 ns between two residues. Our work highlights the importance of other system variations, such as chain length, in modulating the material properties of biocondensates.

## Results

### Broad conformational ensembles visited by ELPs near the LCST

To access the conformational space visited by ELPs, we begin by computing the free energy profiles as a function of the radius of gyration. The latter is a common choice of collective variable describing conformational, i.e coil-globule, transitions for synthetic polymers^29,30^ and disordered proteins^31^. In Fig. 1 (A) and (B) we compute the free energy profiles for the shortest (*n* = 3) and the longest (*n* = 45) sequences, respectively. We choose the temperature range between 280 K and 325 K which includes the expected LCST window (298 K - 310 K)^32^. For both chain lengths, we observe that more compact conformations are favoured as the temperature of the systems increases which is consistent with the LCST phase behavior of ELPs. The free energy profiles for the shortest sequence are nearly flat with a small preference toward *R*_g_ values between 1 and 1.1 nm at low temperatures and favouring *R*_g_ = 0.8 nm at *T* = 325 K. The energy barriers separating these conformations are of an order of 1 kJ/mol, resulting in broad conformational ensembles as was reported earlier^33^. For the longest sequence, the free energy profiles display a two-state behavior with a modest energy barrier ∼10 kJ/mol separating conformations larger than *R*_g_ = 2.2 nm. At *T* = 280 K, 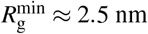 nm, while at *T* = 325 K broad range of conformations with *R*_g_ *<* 2.2 nm is sampled with two minima at 2.1 and 1.8 nm, respectively. Thus the compact state, i.e globule, does not correspond to a single *R*_g_ value, but rather to a set of conformations with a relatively wide range (∼0.4 nm) in *R*_g_. Similar observations were made for a synthetic polymer, which also exhibit the LCST phase behavior^29^. To ensure that the *R*_g_ range is sufficient to sample the free energy profile for *n* = 45, we extend our calculations to *R*_g_ = 6 nm (see SI, Fig. S1 (A)), which confirm that the minima depicted in Fig.1 (B) are the global ones. The free energy profiles for ELPs with *n* = 5 and 12 are shown in Fig. S1 (B) and (C) in the SI.

**FIG. 1.**
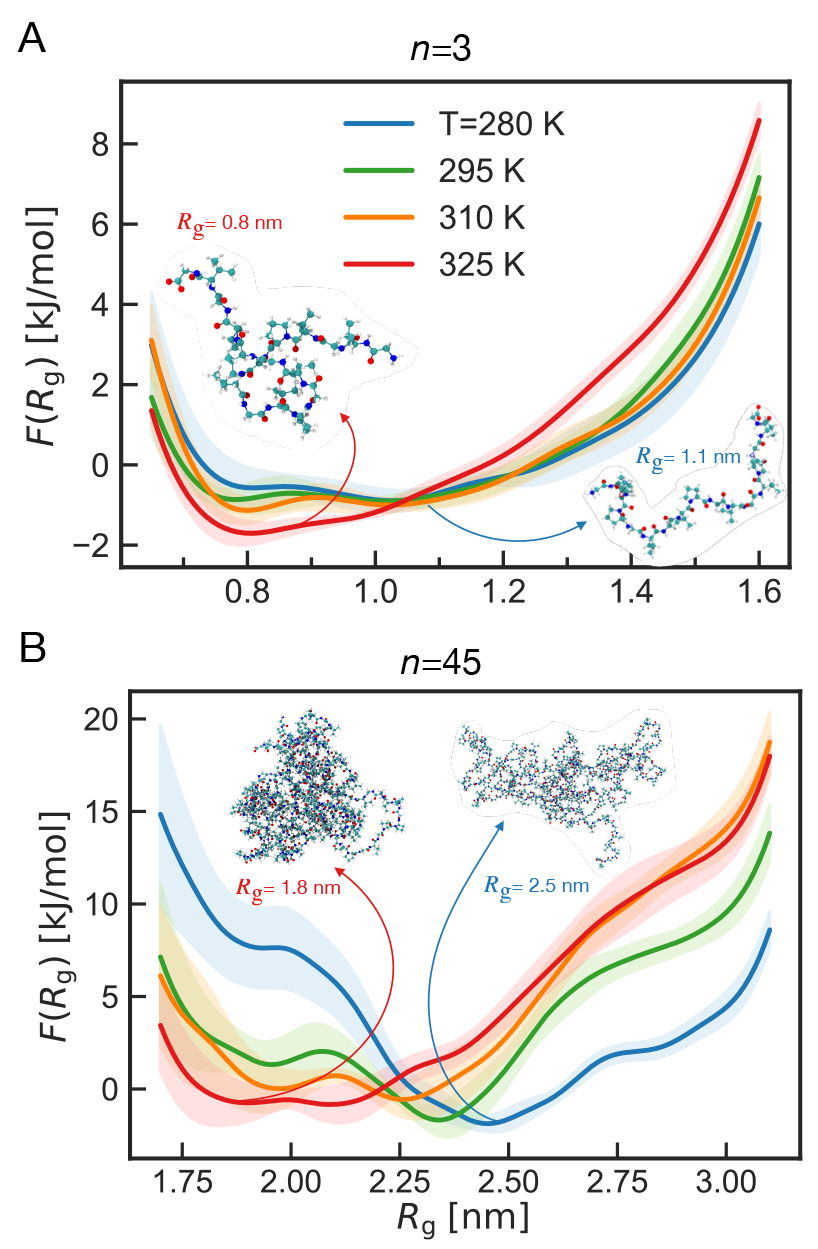
Average free energy profiles of an ELP with *n* = 3 (A) and *n* = 45 (B) pentamers, respectively, vs the radius of gyration below, at, and above the LCST.

### Structural properties of ELPs exhibit strong chain length dependence

To investigate conformational properties of ELPs below and above the LCST, we perform three independent simulations in the *NVT* ensemble using conformations corresponding to the minimum of the free energy profiles for a given (*n, T*) as starting configurations (see Materials and Methods for details). To quantify the conformations visited by ELPs we compute several quantities from polymer physics that help to distinguish between coil and globule states. In Fig. 2 (A) and (B) we compute the root-mean-square internal distances *R*_*i j*_ between residues as a function of their separation |*i* − *j*| along the sequence, for *T* = 280 K and 325 K, respectively. At small separation distances |*i* − *j*| ≤4 the curves are indistinguishable as imposed by chain connectivity. However, at larger separations along the sequence, the scaling of *R*_*i j*_ ∝ |*i* − *j*| ^*ν*^ is qualitatively different among ELPs considered. The sequences with three and five pentamers exhibit coil-like behavior, where the former can be described by a self-avoiding walk (SAW) with the Flory exponent *ν* ≈ 0.59^34^, and the latter is closer to an ideal chain configuration with *ν* = 0.5^35^. Polypeptides with *n* = 12 and 45 repeat units exhibit globular-like behavior, where *R*_*i j*_ reaches a plateau value which was also noted for other disordered protein, polypetides, and synthetic polymers in globular conformations^16,36,37^. The plateau value demonstrates that the actual distance between two residues is independent of |*i* − *j*| for large enough separation along the sequence and equals to the size of a globule, which is 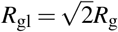 ^38^. That explains the difference in the plateau values for *n* = 12 and 45 pentamers, as the size of the latter is always greater.

**FIG. 2.**
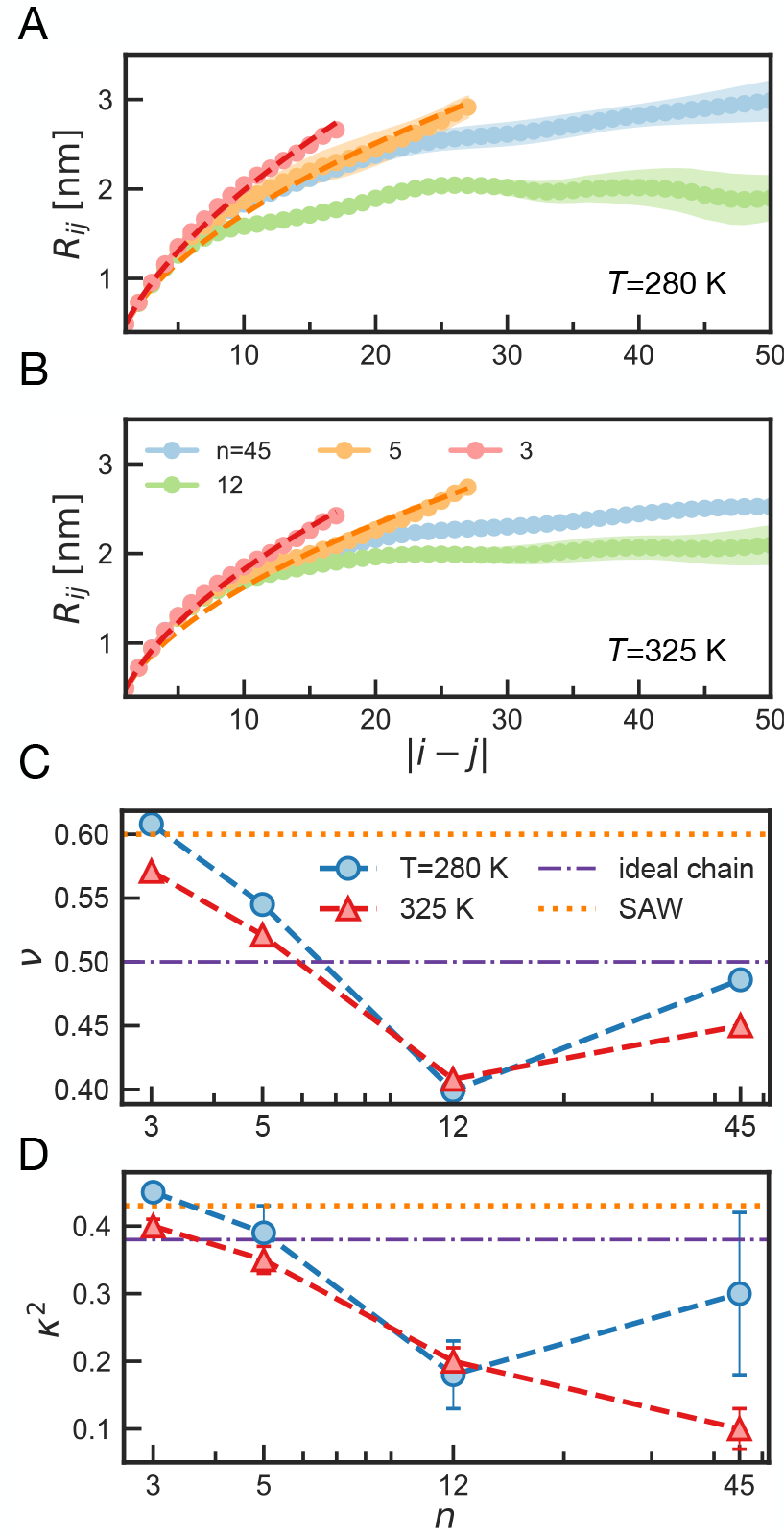
Structural properties of ELPs. (A) and (B) root-mean-square distances *R*_*i j*_ between residues *i* and *j* as a function the distance along the sequence |*i* − *j*| for four ELPs investigated at two temperatures *T* = 280 K and *T* = 325 K, respectively. Dashed curves correspond to plots *R*_*i j*_ = *b* |*i* − *j*| ^*ν*^, where *b* = 0.49 nm is the distance for |*i* − *j*| = 1. (C) and (D) the Flory exponent *ν* and the relative shape anisotropy *κ*^2^ as a function of number of pentamers *n* in a ELP sequence for two temperatures investigated. Dotted and dash-dotted lines correspond to the self-avoiding walk (SAW) and an ideal chain limits, respectively.

We summarize the extracted *ν* values in Fig. 2 (C). We note that *ν* values for sequences with *n* = 12 and 45 are rather approximate, as chain dimensions are too small to ensure a large scale fractal nature^31^. We also characterize the shape of an ELP by computing the relative shape anisotropy parameter *κ*^2^, which attains a value *κ*^2^ = 0 for a fully isotropic shape, e.g. sphere, and *κ*^2^ = 1 for a perfect rod (see SI for details). For a randomly coiling chain in three dimensions, a value of *κ*^2^ = 0.38 is expected in the limit of *N* → ∞.^39^, while for a SAW *κ*^2^ = 0.43.^40^ In accord with the Flory exponent values, ELP with *n* = 3 and *n* = 5 adopt SAW and ideal chain configurations, respectively. While the longer chains display *κ*^2^ values around 0.2 which is closer to spherical globules. We note that a large uncertainty for *κ*^2^ at *n* = 45 at *T* = 280 K originates from chain extension in one trajectory (see Fig. S2, SI) in accordance with the shallow nature of the free energy profiles shown in Fig. 1 (B). Thus, we show that conformational ensembles visited by ELPs are qualitatively different among short and long sequences, and temperature has only a mild effect on conformational ensembles with more compact states visited at higher *T* for all sequences.

### Internal hydrogen bonds define coil *vs* globule states in ELPs

To get a microscopic understanding of why short ELP sequences adopt coil-like conformations while the longer ones (*n* > 5) remain globules, we investigate the formation of intrapeptide hydrogen bonds (iHBs), as it is known that it is crucial for macromolecules that exhibit the LCST phase behavior^37^. In Fig. 3 (A) and (B) we show the number of iHBs per residue, *ID*_res_, as a function of simulation time for sequences with *n* = 3 and *n* = 45 pentamers, respectively. There is a striking difference in the lifetime of iHBs between these two chain lengths. Residues in the short chain engage in iHBs only temporarily, which is consistent with the dynamical nature of ELPs for this chain length^33^. Amino acids belonging to the long chain can form iHBs that last almost a whole trajectory (> 900 ns). Indeed, the lifetime of iHBs, 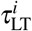, show strong chain length dependence (see Fig. 3 (C)), where 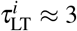 ns and 42 ns for *n* = 3 and 45, respectively. The lifetime 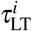 also depends on the temperature, where hydrogen bonds have shorter timescales at higher *T* for all chain lengths considered. However, *T* has only a mild effect on 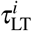, consistent with results from the structure calculations (Fig. 2). We also compute the lifetime of the polypeptide-water hydrogen bonds 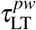 (see Fig.S3 in the SI), which exhibit similar to 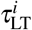 behavior with one noticeable difference: 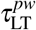 is predominantly in the sub-nanosecond range with the maximum value of 1.3±0.4 ns for *n* = 45 at *T* =280 K. Thus, we can unambiguously conclude that the timescales of the intrapeptide hydrogen bonds define the conformational destiny of an ELP.

**FIG. 3.**
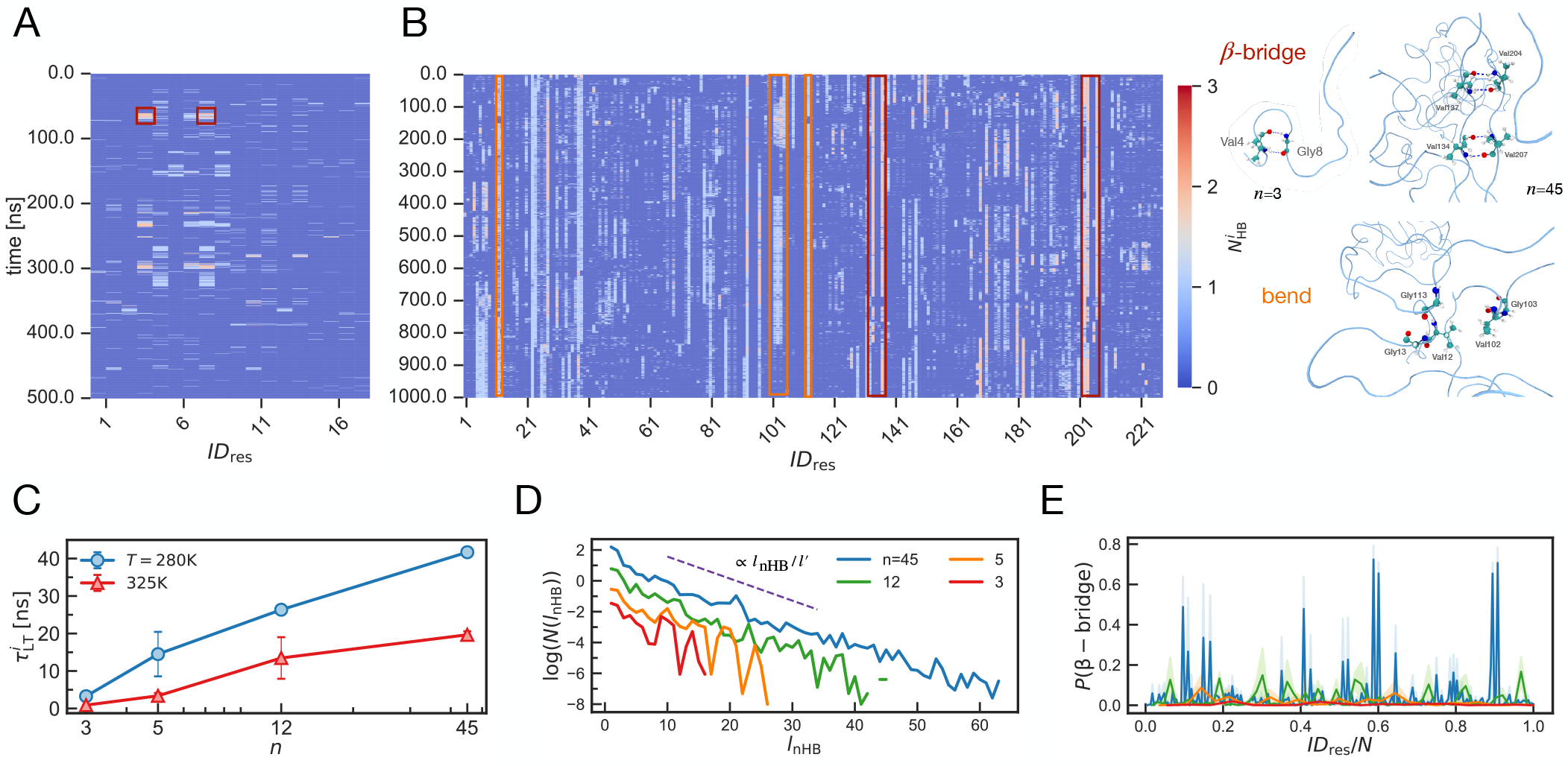
Intrapeptide hydrogen bonds: their lifetime and distribution. (A) and (B) spatial-temporal evolution of the number of intrapeptide hydrogen bonds (iHB) 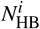 per residue at *T* = 280 K for the sequences with *n* = 3 and *n* = 45 pentamers, respectively. MD snapshots on the right correspond to HBs highlighted by red (*β*-bridge) and orange (bend) rectangles. (C) Lifetime of the intrapeptide hydrogen bonds 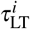 as a function of the number of pentamers *n* below and above the LCST temperature. (D) The average number of the consecutive non-hydrogen bonded (nHB) residues *N*(*l*_nHB_) at *T* = 280 K for four sequence lengths investigated, where *l*^*′*^ = 7 is the mean sequence length between two hydrogen-bonded residues. (E) The average probability of *β*-bridge formation per residue at *T* = 280 K for four sequences investigated.

To quantify the separation distance between two residues that form an iHB, In Fig. 3 (D), we compute the average number *N* of consecutive nonhydrogen bonded (nHB) residues *l*_nHB_^41^. It follows an exponential decay with the characteristic length scale *l*^*′*^ = 7 ± 1 residues at all *n* considered. However, the occurrence of subsequences of length *l*^*′*^ is extremely small (⪅ 0.1) for chains with *n* = 3 and 5 forming coils. This is in contrast to longer chains (*n* = 12 and 45), where *N*(*l*^*′*^) > 0.5 which populate globule-like states.

We further investigate the occurrence of the long-lived iHB shown in Fig. 3 (B). We find that these iHBs often correspond to residues that form either a *β*-bridge or a bend/HBturn as a type of secondary structure. We show examples of *β*-bridge or bend forming residues by red and orange rectangles in Fig 3 (A, B) and the corresponding MD configurations on the right-hand side. For example, two *β*-bridges formed by Val134-Val207 and Val137-Val204 (up to four hydrogen bonds in total) exist for more than 600 ns. By restricting the mobility of these four amino acids, the residues between them still retain high conformational entropy by forming coils, bends, and turns. In Fig. 4 (E), we show the probability of the *β*-bridge formation as a function of *ID*_res_ scaled by the total chain length *N* = 5*n* + 3. Indeed, for *n* = 45 and *n* = 12, this type of secondary structure occurs every 10 to 20 residues. For shorter sequences these conformations are rarely seen, which is due to their short length of *N* = 18 and 28 residues for *n* = 3 and 5, explaining their coil-like conformations.

**FIG. 4.**
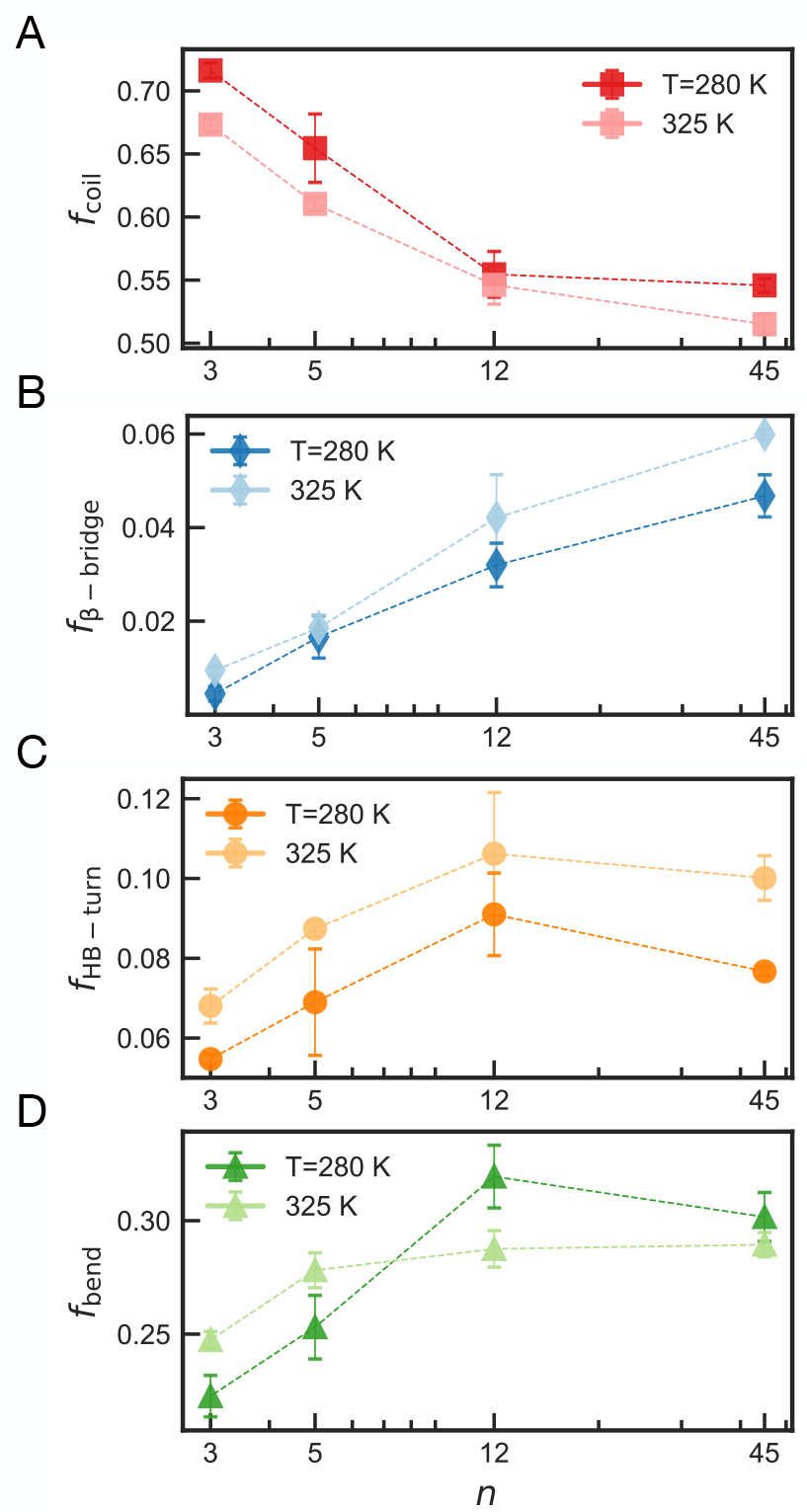
Fraction of residues attributed to the following secondary structure motifs: (A) coil *f*_coil_, (B) *β*-bridge *f*_*β* −bridge_, (C) hydrogen-bonded turn *f*_HB−turn_, and (D) bend *f*_bend_ as a function of the number of pentamers *n* for temperatures below (280 K) and above (325 K) the LCST.

### Number of pentamers controls the extend of disorder in ELPs

To better quantify the difference in secondary structure formed by ELPs of different length *n*, in Fig. 4 we compute the fraction of residues attributed to coils (A), *β* −bridges (B), hydrogen-bonded turns (C), and bends (D). Overall, more than half of residues belong to coil states, which is in agreement with the disordered nature of ELPs. However, the exact fraction varies significantly with the number of pentamers *n*, with shorter chains reaching up to 70 % of the sequence belonging to coils while being around 50 % for the longest ones. Thus, amino acids belonging to longer sequences participate more in ordered motifs such as *β* −bridges and hydrogen-bonded turns or bends. This tendency remains at both temperatures below and above the LCST, where the ELPs are more ordered above the LCST. Thus, we find that both temperature and chain length affect the structural properties of ELPs. However, the temperature provide a slight bias toward more ordered secondary structure motifs. The chain length, in contrast, qualitatively changes the conformational ensembles visited by ELPs and can be seen as a parameter that can control the extension of the ratio between disordered and ordered elements in chain.

## Discussion

Control over aggregate properties of biocondensates is essential for both combating pathological assemblies in cells and for application-specific designs of biomaterials. For example, elastin-based hydrogels with fast gelation time are well-suited for drug delivery applications^42^, while materials with longer gelation time (∼days) are desired for tissue engineering^43^. It was shown that for disordered proteins, single-chain properties such as the chain size correlate well with the propensity of the sequence to undergo phase separation^20–22^.

By acknowledging this similarity between inter- and intra-chain interactions, here we studied ELPs characterized by the LCST phase behavior at the single chain level. We varied the number of pentameric repeats from 3 to 45 covering a wide range of chain lengths. We restricted our study to temperatures between 280 and 325 K, which includes the experimentally observed critical temperature for the sequences investigated, while remaining suitable for possible applications. For the first time, the free energy profiles of ELPs were computed, and we report that all sequences exhibit compaction in size with temperature increase, consistent with the thermoresponsive nature of ELPs (Fig. 1, and Fig. S1 in the SI). However, when we characterized the conformational ensembles of ELPs employing descriptors borrowed from polymer physics such as the Flory exponent, and the shape of the polypeptide chain, we observed that short sequences predominantly visit coil-like states, while the longer ones remain globules (see Fig. 2). We rationalized our observations by computing the distribution and timescales of the intrapeptide hydrogen bonds. Our calculations revealed that short peptides form intrapeptide hydrogen bonds only temporarily (~a few ns) highlighting their dynamic nature consistent with former numerical and experimental studies for peptides of similar length^25,27,33^ shown in Fig 3 (A). In contrast, longer polypeptides engage in long-lived internal hydrogen bonds that can last hundreds of nanoseconds, effectively restricting the conformational space to more compact, e.g. globule, conformations (see Fig 3 (B)). We attribute this striking difference between peptides of different lengths to the effective stiffness of a chain. Short chains effectively appear as more rigid, as the number of conformations allowed by compact, e.g. globule, states is low, which in turn reduces the conformational entropy of the chain. In longer sequences, by restricting mobility of a few residues that form *β*-bridges, the subsequence in between these residues is allowed to sample a large conformational space with interconverting coils, HB-turns and bends secondary structure motifs. We showed that the sequence length between two non-hydrogen bonded amino acids is on average seven residues (Fig 3 (D)), which can be interpreted as the Kuhn length - the length scale that quantifies the stiffness of a polymer chain. Thus, the short sequences can be interpreted as oligomers with just three to four monomers, while the longer sequences resemble polymers with tens of freely-joined segments. Similar observations were reported for synthetic polymers where polystyrene oligomers remain in coil conformations even when dissolved in poor solvents^30^. This difference in ELP conformations is further explained through the secondary structure formation shown in Fig. 4. The amino acids in short sequences are mostly assigned to coils, while residues in polypeptides have a higher propensity to form ordered structures such as *β*-bridges, HB-turns, and bends. Thus, the length of the sequence controls the degree of disorder in ELPs.

We propose, that this ability to form long-lived internal hydrogen bonds in polypeptides might help to rationalize the recent experimental findings, where mini-elastin capped with hydrophobic domains (HDs) of 30-42 residues formed liquid droplets while doubling the length of HDs resulted in the formation of solid-like condensates^28^. The sequence lengths of HDs used in experiments agree remarkably well with our numerical findings, where we observe the formation of globule states for ELPs with 63 residues and larger, and coils states for shorted sequences. We further generalize our findings and speculate that the aggregate state of elastin-based materials might depend on the length of their hydrophobic domains. The liquid droplets are formed by polypeptides with short HDs, while the assembly of sequences with long HDs results in solid aggregates unifying former numerical and experimental viewpoints. Naturally, the sequence composition has a profound effect on the conformational properties of proteins and polypeptides; however, the length of the sequence provides an additional powerful variable for finely tuning material properties, which previously did not receive much attention in the literature.

## Materials and Methods

We performed explicit solvent, all-atom molecular dynamics (MD) simulations employing the AMBER99SB-ILDN force field^44^ and the TIP4P-D water model.^45^ This combination of the force field and water model was shown to be suitable for modeling disordered proteins^46^ including thermoresponsive behavior of short ELPs^33^. Additionally, we demonstrated that the TIP4P-D water model reproduces well thermodynamic, dynamic, and dielectric properties of liquid water in a wide temperature range including 280 K to 325 K which is of interest here^47^.

We considered four ELPs sequences with number of VPGVG pentamers *n* equaled to 3, 5, 11, and 45. We additionally added GVG unit to the beginning of each peptide to mimic experimentally available sequences, thus total length *N* of the sequences investigated was 18, 28, 63, and 228 residues. The initial configurations were assembled using the Avogadro molecular builder (ver. 1.20)^48^. In our modeling, we consider neutral pH values which imply that the N and C-termini of the peptide are charged (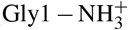 and GlyN − COO^−^, respectively). To mimic experimental conditions, we added 100 mM of NaCl to the systems. The ELPs were solvated in a cubic box with the initial sizes ranging from *l*_box_ = 6 nm to *l*_box_ = 15 nm. This box size ensures that the chains do not interact with their periodic images as the box sizes exceeds the chain size (the radius of gyrations) at least three times. All simulations were conducted using the software package GROMACS 2020.2^49,50^.

For each system, we began with a preliminary run in the *NpT* ensemble at *T* =298 K and at a pressure of 1 bar where a peptide visited a set of confirmations. The length of these simulations varied with the peptide length *n* and reached 2000 ns for the longest sequence considered (*n* = 45). These simulations allowed us to identify the peptide’s size range for the subsequent free energy profile calculations.

To construct the free energy profiles, we used the Umbrella Sampling (US) method with the radius of gyration *R*_g_ as the collective variable which was fixed over a simulation run, e.g. sampling window, by a harmonic potential as 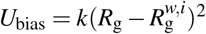, where *k* is the force constant, 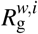 is the desired value for the *i*th window. We computed *R*_g_ using positions of *C*_*α*_ atoms only. The window spacing of 0.05 nm was adopted. We varied values of *k* to ensure the sufficient overlap between neighboring windows. For the longest chain with *n* = 45 pentameric units *k* = 6000 kJ/(mol·nm^2^), while for the shortest one *k*(*n* = 3) = 4000 kJ/(mol·nm^2^). Each window was simulated for 15 ns. We conducted five independent US calculations at each temperature and chain length to compute the average free energy profile for this state point. The weighted histogram analysis method (WHAM) was employed to derive the free energy profile curves *F*(*R*_g_). These calculations were carried out using the open-source, community-developed PLUMED library (ver. 2.8.[1])^51,52^.

To compute the structural and dynamical properties of ELPs below (280 K) and below (325 K) the LCST, we conducted three independent simulations on the *NVT* ensemble where we used the chain configuration with the radius of gyration corresponding to the minimum in the free energy profile at a given state point. The simulation length was set to 500 ns which exceeds at least by an order of magnitude the end-to-end relaxation time *τ*_R_ of ELPs investigated except systems with *n* = 45 at *T* =280 K where it was increased to 1000 ns as *τ*_R_ ≈ 80 ns for this system.

We analyzed trajectories using in-house scripts as well as GROMACS analysis tools such as the implementation of the DSSP algorithm^53,54^ for evaluating peptide secondary structure, the number of hydrogen bonds, a python library MDTraj^55^ for computing pairwise distances, the gyration tensor.

## Author Contributions

T.I.M and J.-L. B. designed the research, T.I.M performed simulations and subsequent analysis; T.I.M wrote the first draft; all authors contributed to the preparation of the manuscript.

## Data availability

The data that support the findings of this study are available from the corresponding author upon reasonable request.

## Acknowledgments

This work was performed using HPC resources (GPU-accelerated partitions of the Jean Zay supercomputer) from GENCI–IDRIS (Grant 2021-2023 - A0100712464). J.-L.B. and N.A.G. acknowledges International Research Project “Statistical Physics of Materials” (CNRS-CONICET).

