## Supporting Information for "Sequence length controls coil-to-globule transition in elastin-like polypeptides"

### Free energy profiles

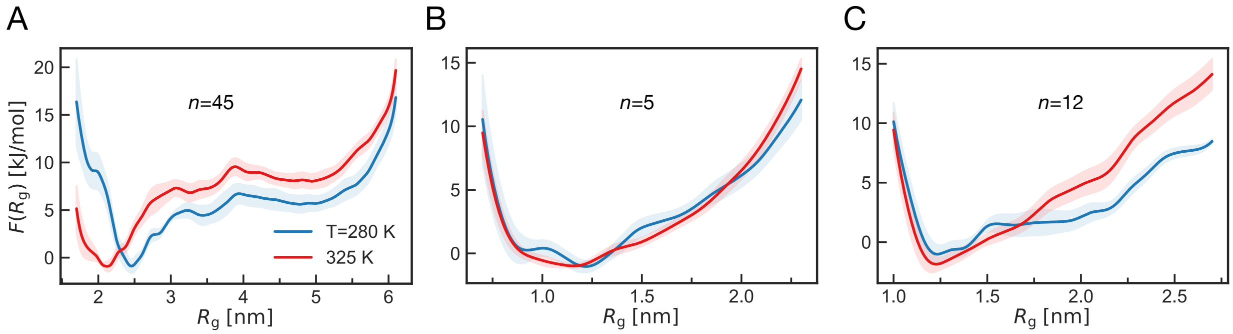


Figure 1: (A), (B), and(C): Free energy profiles of the sequence with $n = 45$, 5 and 12 pentamers, respectively, at temperatures below ($T = 280$ K) and above ($T = 325$ K) the LCST.

### Calculation and scaling of the internal distances

We compute the root-mean-square internal distances $R_{ij}$ between $i$ and $j$ residues using the positions of all atoms belonging to these residues as:

$$R_{ij} = \frac{1}{M_{ij}} \sum_{k \in i} \sum_{l \in j} |\mathbf{r}_k - \mathbf{r}_l| \tag{1}$$

where $\mathbf{r}_k$ and $\mathbf{r}_l$ is the positions of $k$ and $l$ atoms that belong to $i$ and $j$ residues, respectively, and $M_{ij}$ is the number of unique atom pairs between $i$ and $j$ residues. We find that the mean distance between two consecutive amino acids $|i-j| = 1$ is equal to $b = 0.49$ nm and neither depends on the sequence length nor temperature, as was noted earlier for other disordered proteins.[1,2] We use this value as a prefactor in our fitting procedure $b|i-j|^{\nu}$.

### Calculation of the radius of gyration and the relative shape anisotropy

To quantify the size and the shape of ELPs, we define the gyration tensor $\mathbf{G}$ as

$$G_{\alpha\beta} = \frac{1}{N_a}\sum_{i=1}^{N_a} \Delta r_\alpha^i \Delta r_\beta^i \tag{2}$$

where $N_a$ is the number of atoms in a chain, $\Delta r_\alpha^i$ is the position of atom $i$ relative to the chain center of mass, while $\alpha$ and $\beta$ are the components in the Cartesian $x$, $y$, and $z$ directions. The chain radius of gyration is defined as $R_\mathrm{g}^2 = G_{xx} + G_{yy} + G_{zz}$. Note, that for the Umbrella Sampling calculations, we used positions of $C_\alpha$ atoms only. Using the instantaneous values of the eigenvalues of the gyration tensor $\mathbf{G}$: $\lambda_1$, $\lambda_2$, $\lambda_3$, we compute the relative shape anisotropy $\kappa^2$ of an ELP as

$$\kappa^2 = 1 - 3\left\langle \frac{\lambda_1^2\lambda_2^2 + \lambda_1^2\lambda_3^2 + \lambda_2^2\lambda_3^2}{(\lambda_1^2 + \lambda_2^2 + \lambda_3^2)^2} \right\rangle \tag{3}$$

where the angular brackets denote averaging over a simulation trajectory.

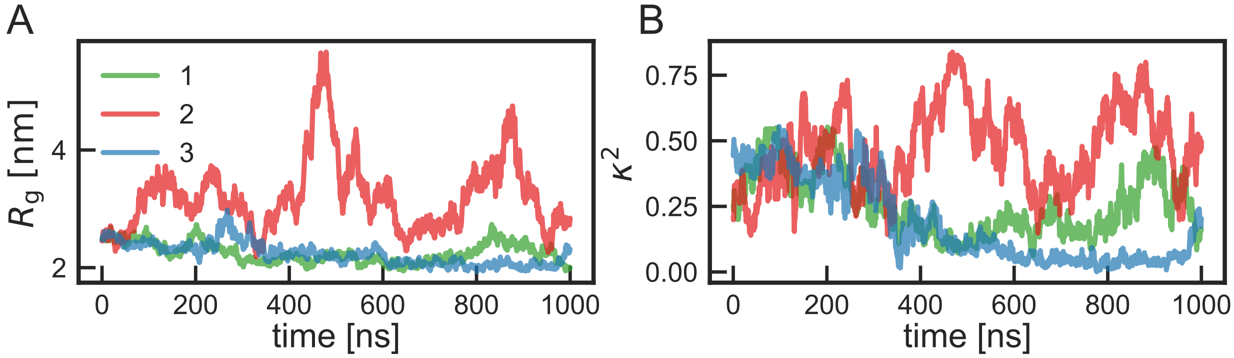


Figure 2: (A) and (B): the temporal evolution of the radius of gyration $R_\mathrm{g}$ and the relative shape anisotropy $\kappa^2$, respectively, for $n = 45$ at $T = 280$ K for three independent simulations.

#### Calculation of the lifetime of hydrogen bonds

We used the standard geometrical definition of a hydrogen bond (HB), where a HB exists if the donor-acceptor distance is $\leq 0.35$ nm and the acceptor-donor-hydrogen angle is $\leq 30°$. To estimate the mean lifetime of HBs (intrapeptide and peptide-water), we first computed the existence matrix $S_{ij}(t)$ of hydrogen bonds at each timestep of a trajectory using *gmx hbond -hbm* function implemented in GROMACS. The value of $S_{ij}(t)$ is one if a HB is formed between residues $i$ and $j$ at time $t$, and zero otherwise. Then we compute the average over all existing HBs autocorrelation function (ACF) as

$$C(t) = \frac{\sum_{ij} S_{ij}(t_0) S_{ij}(t + t_0)}{S_{ij}(t_0)}. \tag{4}$$

We computed the so-called intermittent ACF, which gives the probability of a HB being intact at time $t$ given that it existed at time $t_0$, i.e. breaking events at the times between $t_0$ and $t$ were not taken into account.[3,4] Finally, we integrated ACFs over this time interval equal to half of a trajectory length subtracting ACF's long-time limit. The values of the lifetime reported in this work are averaged over three independent system realizations.

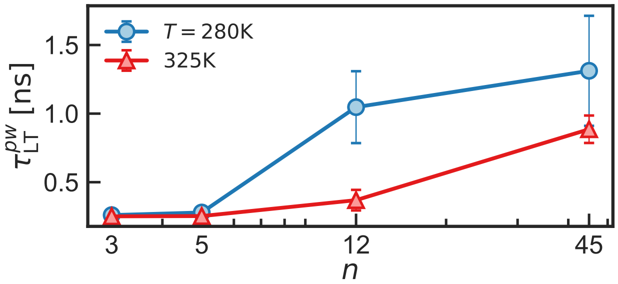


Figure 3: Lifetime of the polypetide-water hydrogen bonds $\tau_{\mathrm{LT}}^{pw}$ as a function of the number of pentamers $n$ below ($T = 280$ K) and above ($T = 325$ K) the LCST.
